# Human *Chrysomya bezziana* myiasis: A systematic review

**DOI:** 10.1101/610121

**Authors:** Xianyi Zhou, Dzinkambani Moffat Kambalame, Sitong Zhou, Xiang Guo, Dan Xia, Yemei Yang, Rangke Wu, Juan Luo, Fenglong Jia, Mingchi Yuen, Yuehua Xu, Geyang Dai, Li Li, Tian Xie, Santhosh Puthiyakunnon, Wenxia Wei, Lixian Xie, Siting Liang, Yuqin Feng, Songgen Huang, Yongxuan Hu, Qianzhen Mo, Rongjia Mai, Xiaoqing Zhang, Philip Spradbery, Xiaohong Zhou

**Affiliations:** Department of Dermatology, The 3rd Affiliated Hospital, Southern Medical University, Guangzhou 510630, Guangdong, China; Department of Pathogen Biology, Key Laboratory of Prevention and Control for Emerging Infectious Diseases of Guangdong Higher Institutes, Guangdong Provincial Key Laboratory of Tropical Disease Research, School of Public Health, Southern Medical University, Guangzhou 510515, Guangdong, China; Department of Dermatology, Shenzhen Hospital, Southern Medical University, Shenzhen 518100, Guangdong, China; The School of Foreign Studies, Southern Medical University, Guangzhou 510515, Guangdong, China; Institute of Entomology, Life Sciences School, Sun Yat-sen University, Guangzhou 510275, Guangdong, China; Education Technique Center, Southern Medical University, Guangzhou 510515, Guangdong, China; Department of Dermatology, Nanfang Hospital, Southern Medical University, Guangzhou 510515, Guangdong, China; XCS Consulting, Yarralumla, Canberra, ACT 2600, Australia

## Abstract

**Background:** Myiasis due to Old World screw-worm fly, *Chrysomya bezziana*, is an important obligate zoonotic disease in the OIE-list of diseases and is found throughout much of Africa, the Indian subcontinent, southeast and east Asia. *C. bezziana* myiases cause not only morbidity and death to animals and humans, but also economic losses in the livestock industries. Because of the aggressive and destructive nature of this disease in hosts, we initiated this study to provide a comprehensive understanding of human myiasis caused by *C. bezziana*.

**Methods:** We conducted a systematic search of the databases in English (PubMed, Embase and African Index Medicus) and Chinese (CNKI, Wanfang, and Duxiu), and international government online reports to 6^th^ February, 2019, to identify studies concerning *Chrysomya bezziana*. Another ten human cases in China and Papua New Guinea that our team had recorded were also included.

**Results:** We retrieved 1,048 reports from which 202 studies were ultimately eligible for inclusion in the present descriptive analyses. Since the first human case due to *C. bezziana* was reported in 1909, we have summarized 291 cases and found that these cases often occurred in poor hygiene, low socio-economic conditions, old age, and underlying diseases including infections, age-related diseases, and noninfectious chronic diseases. But *C. bezziana* myiasis appears largely neglected as a serious medical or veterinary condition, with human and animal cases only reported in 16 and 24 countries respectively, despite this fly species recorded as present in 44 countries worldwide.

**Conclusion:** Our findings indicate that cryptic myiasis cases due to the obligate parasite, *C. bezziana,* are under-recognized. Through this in-depth study to clarify the knowledge of human *C. bezziana* myiasis including its etiology, clinical features, diagnosis, treatment, epidemiology, prevention and control, we call for more vigilance and awareness of the disease from governments, health authorities, clinicians, veterinary workers, nursing homes, and also the general public.

**Author summary:** *Chrysomya bezziana* larvae are characterized by feeding aggressively on the living tissues and body fluids of the host. The nightmare-like feelings of patients suffering from this myiasis, severe tissues and bones destruction, even death, and enormous economic loss in the livestock industries have been described in the previous reports. But our findings indicate that *C. bezziana* myiases still appear to be under-recognized as a serious medical or veterinary condition throughout the world. Both in China and the world at large, it is probable that *C. bezziana* distribution could be greater than currently reported. For the first time, we have therefore systematically investigated human myiasis caused by *C. bezziana*. Our study provides an opportunity for clinicians and health authorities to gain a better understanding of this disease from its etiology, pathology, clinical features, diagnosis, treatment, epidemiology, prevention and control. In addition, our findings will hopefully engage governments, health staff, veterinary workers, the elderly homes, and also the general public in efforts to recognize, prevent and control such infestations.

## Introduction

> It is doubtful that the mind of man could create a more vile scene than that of worms consuming the live flesh of one’s body. The imagination almost refuses - particularly in this day and age - to conjure up the horrendous pain and outright revulsion that must come to a person infested with a writhing, seething mass of worms steadily tearing and consuming his flesh.
>
> — --C.G. Scruggs [1]

The Old World screw-worm fly *Chrysomya bezziana*, is an obligate parasite, belonging to the order Diptera, family Calliphoridae, and suborder Cyclorrhapha. It is distributed throughout much of southeast Asia, the southern part of east Asia, the Indian subcontinent, Papua New Guinea (PNG), the Middle East and tropical and subtropical Africa [2-3]. Myiasis due to *C. bezziana* is one among 117 OIE-listed diseases (World Organisation for Animal Health, http://www.oie.int/animal-health-in-the-world/oie-listed-diseases-2019/). The first case of *C. bezziana* myiasis was reported in cattle in 1909 [4]. Since then, sporadic cases and even major outbreaks of myiases have been reported globally in animals [5]. Human myiasis caused by *C. bezziana* was firstly reported in 1909 in India [6], and to the present time, at least 291 cases have been reported worldwide in humans (S1 Table). *C. bezziana* larvae are characterized by feeding on the living tissues and body fluids of the host, so once the infestation occurs, it can cause serious destroy to the tissues even bones of the host, and if vital organs are involved, death may occur [6]. Not only does *C. bezziana* cause morbidity and death to animals and humans, but also economic losses in livestock industries [5,7]. Such terrible suffering from this myiasis was described as a nightmare in the previous studies [1,7,9]. Despite the aggressive and destructive nature of this disease in hosts, its epidemiology, etiology, pathology, clinical features, diagnosis, treatment, prevention, and control in humans remain inadequately characterized. Therefore, we initiated this study to comprehensively investigate human myiasis due to *C. bezziana*.

## Materials and methods

### Search strategy and selection criteria

We conducted a systematic research of the databases relating to *C. bezziana* myiasis in humans in English and Chinese up to 6^th^ February, 2019, including PubMed (https://www.ncbi.nlm.nih.gov/pubmed/), Embase (https://www.elsevier.com/solutions/embase-biomedical-research), the African Index Medicus (AIM, http://indexmedicus.afro.who.int/), the China National Knowledge Infrastructure Databases (CNKI, http://www.cnki.net), Duxiu Scholar (http://www.duxiu.com/), and Wanfang (http://g.wanfangdata.com.cn/). We used the following search terms: (chrysomya bezziana) OR (c. AND bezziana) OR (c. bezziana) OR (chrysomya) OR (chrysomya AND bezziana AND villeneuve) OR (chrysomya AND bezziana AND vill) OR (old world screwworm) OR (old AND world AND screwworm) (in English), 蛆症金蝇 or 倍氏金蝇 or 白氏金蝇 or 旧世界螺旋虫(in Chinese). As well as the government online reports in the world were retrieved, for example, WHO/TDR (http://www.who.int/tdr/en/), Centers for Disease Control and Prevention in USA (https://www.cdc.gov/), and National Health Commission in China (http://www.moh.gov.cn/zhuz/index.shtml), etc., but the eligible government documents utilized for further study only verified from Centre for Health Protection, Department of Health, the Government of the Hong Kong Special Administrative Region (CHP, http://www.chp.gov.hk/). The websites of Invasive Species Compendia from the Commonwealth Agricultural Bureau International (CABI) (https://www.cabi.org/ISC/) were also searched. All studies concerning etiology, pathology, clinical features and epidemiology of *C. bezziana* myiasis, including case reports, case series, reviews, cross-sectional and cohort studies were retrieved. Articles were screened by title and abstract and full-text articles were then assessed for eligibility. The primary outcome was to exclude key words 'new world screwworm'. No restrictions on date of publication or country of origin were applied. For those reports with available full texts, references were checked for potential reports. A pre-piloted electronic form designed in Microsoft Access was used to extract data from the reports. After duplicates were removed, all the references were checked and in those not specified by species identification of *C. bezziana* by authors were excluded. Another ten human cases of *C. bezziana* in China (1) and Papua New Guinea (9) that our team have recorded were also included. The authors (DMK, XG, STZ, LXX, and XG, YMX, WXW, STL), separately conducted the search and selection of the databases in English and Chinese and, if any conflicts arose, they resolved them by consulting the senior authors (XHZ, SP, XYZ, FLJ and MZY). We also consulted some of the core experts in the field when needed. Datasets of maps were download from the Natural Earth (Free vector and raster map data @ naturalearthdata.com.), and maps of geographical distribution of *C. bezziana* as well as the disease in the world and in China were conducted using Adobe Illustrator CC 2017.

## Results and discussion

### Study eligibility results

Our PubMed, Embase, and African Index Medicus search in English identified 898 records, of which 199 remained for eligibility, until duplicates were removed, title, abstract and full text screening, in which specified identification of *C. bezziana*. Among them, 4 human cases reported in Algeria, northern Africa [9], Turkey [10], Spain [11], and Mexico [12] were excluded in this analysis due to the doubt of expert opinions on species identification. CNKI, Duxiu Scholar, and Wanfang search in Chinese identified 113 records, of which 26 remained. Meanwhile, distinct from other regions worldwide, *C. bezziana* myiasis-associated data are also available from government documents CDWs (Communicable Diseases Watch) in Hong Kong, 37 eligible records documented by the government of Hong Kong are included. Additionally, we traced related studies from the references of all the included literature as far as possible, as well as the recommendation of the experts, and 44 of them were chosen to enrich the records. Finally, 202 studies were included for descriptive analyses in this review. The PRISMA flowchart was provided in Fig S1.

### Etiology and pathology

The Old World screw-worm fly, *C. bezziana* is one of the most important obligate zoonotic myiasis fly species in tropical and subtropical countries. Its main hosts are cattle, pigs, sheep, dogs, water buffaloes, goats, horses and donkeys as well as native wildlife, while it also invades human hosts [13]. *C. bezziana* infests hosts when eggs are laid by the gravid female, leading to infestations at the edge of a wound, on mucous membranes, at natural body openings or they may cause indirect infestations if a host has close contact with another infested host. Frequently, infestations occur in new-born animals via the umbilical cord [2].

Adult flies of *C. bezziana* are able to mate (Fig 1A) a few days after their emergence (Fig 1F): a male is able to inseminate a female within a day of its emergence, while a female becomes receptive after three days [14]. Females usually mate once but a male is capable of mating with several females during its life time [2]. Three to four days post mating (dpm), females start looking for suitable hosts to lay their eggs. It normally takes a female about 4-11 dpm for the first oviposition to occur [2,15]. Once a host is found, the female lays its eggs on the edge of the open wound (Fig 1B) or at the opening of body cavities such as mouth, ears, eyes or the urogenital system. A female lays up to 245 eggs at one time in the form of a whitish mass and can deposit up to three egg masses in its life time at intervals of 5-18 days [14]. The time taken by the female to lay a single egg mass is about four minutes and once deposited on the host, the mass gets firmly attached to the skin so that it cannot be easily wiped or licked off [14]. It is believed that the females are attracted to the oviposition site by the odour of blood and mucous and especially a prior myiasis. Under favorable conditions, it usually takes 18 to 24 hours at 25 to 30°C for the eggs to hatch into larvae [15], but it can be as short as 10.5 hours at 37°C [14]. The newly hatched larvae undergo three stages of development known as instars. The first-instar larvae migrate to the nearby wound where they feed on blood and wound exudate. After one day, the first-instar larvae moult and develop into the second-instar. During this time, the larvae aggregate and start invading the living tissues with their heads down in the tissues and their posterior end with their spiracles exposed, to enable respiration. One to two days later, the larvae develop into the third-instar during which the strong, highly visible mouth hooks facilitate further destruction of living tissues, accompanied by necrosis and pain at the site of infestation (Fig 1C). The third-instar larvae matures within 7-8 days of the eggs hatching. The larvae begin to leave the wound (Fig 1D) after about 7 days of feeding, with females vacating earlier (day 6-7) than males (day 7-8). The mature larvae leave the host and fall to the ground, burrow into the ground to a depth of about 2.5 cm with their heads downwards, then they turn around so that their heads point upwards and then pupate (Fig 1E) within 1-2 days [15-16]. The pupae develops into the adult fly, the duration of this stage depending on temperature: it only takes 7 days at 28°C or up to 60 days at 10-15°C. Hence, the development of *C. bezziana* from an egg to an adult fly may vary from 15 days to over 60 days, depending on prevailing temperatures. The life span of an adult fly ranges from 16 to 51 days, with an average of 22.4 and 37.5 days for male and female flies, respectively [2,15].

**Fig 1.**
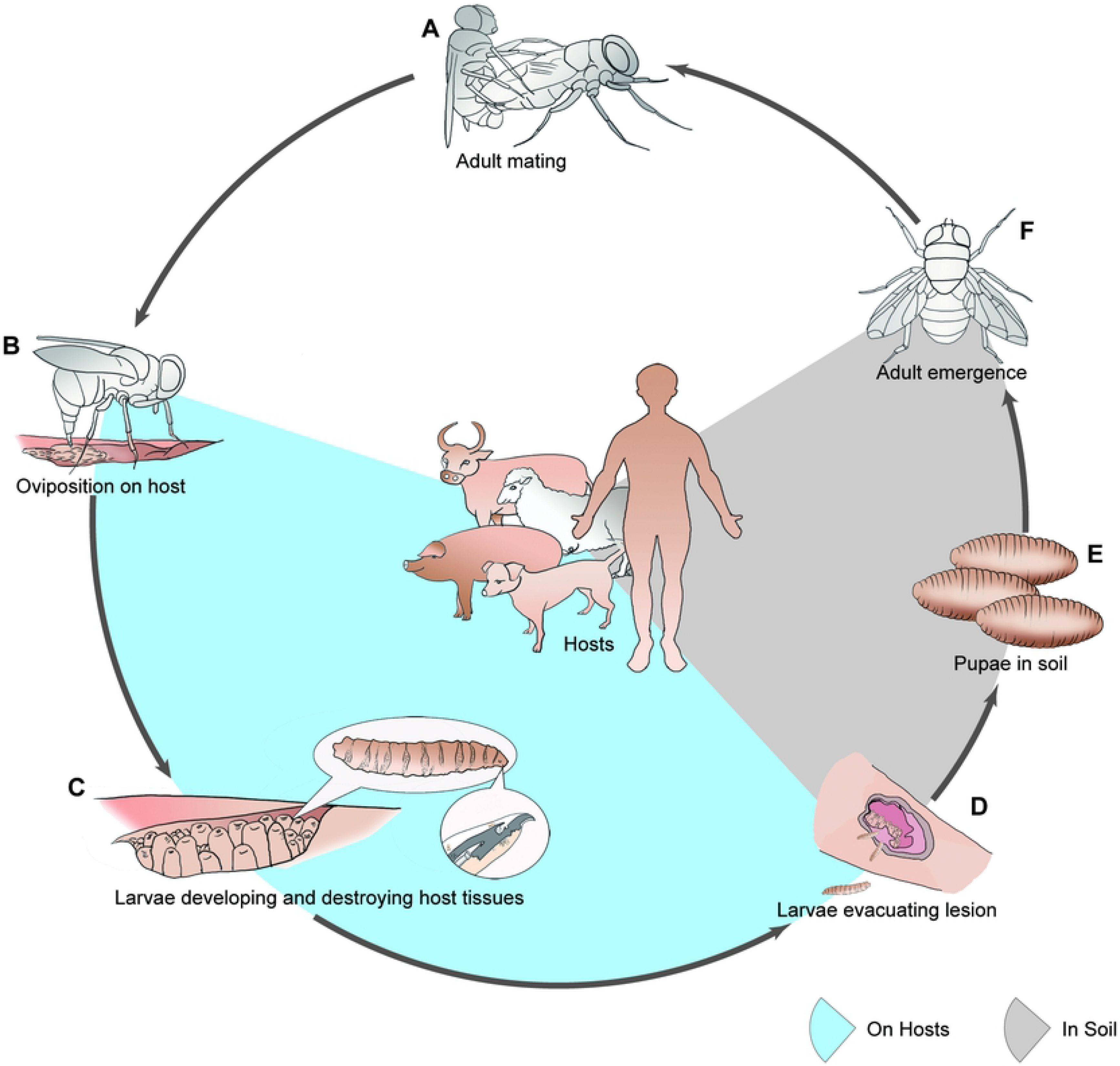
The life cycle of *Chrysomya bezziana*. *Chrysomya bezziana* is an obligate parasite. Cattle are commonly regarded as the principal host of *C. bezziana* [17], and other warm-blooded animal are involved, including native wildlife and domesticated animals, as well humans [13]. (A) Adult flies of *C. bezziana* are able to mate a few days after their emergence. (B) The gravid females lay their eggs on the edge of an open wound of the host. (C) After hatching from the eggs, the larvae undergo three stages of development known as instars through molting, feed on blood, wound exudates and the living tissues of the host, leading to the destruction of the living tissues. (D) After about 7 days feeding, the mature third-instar larvae begin to leave the wound of the host and fall to the ground, burrow into the ground and pupate (E) within 1-2 days. (F) The pupae develop into adult flies.

The gross pathological changes correspond to the larval developmental stages of *C. bezziana* as they feed on the tissues of the host. The larvae can destroy living tissues and cause large ulcerative lesions with bleeding, pain, a serosanguinous purulent discharge, and even fistula. Furthermore, the larvae can cause massive tissue destruction and necrosis, and secondary infections. Consequently, systematic manifestations may occur including fever, chills, weight loss, swollen, and inflammation. The larvae can even invade and destroy bones, nasal sinuses, orbital cavities, hard palate, eyeballs, hearing apparatus and teeth (S1 Table). The aggressive invasion of *C. bezziana* larvae, associated with functional impairment and necrosis, may lead to serious complications such as debility, limb amputation [18-21], eye completely destroyed [6,22-28], and even death if prompt and rapid diagnosis and treatment is unavailable [6]. Recovery can occur when fibrous granulation tissue grows beneath the infested areas. Although the uncomplicated lesions will resolve, the initial massive infestation and/or further strikes on an already infested wound can lead to massive soft tissue destruction and wound extension [29].

### Clinical features

Infestations by *C. bezziana* commonly manifest as wound and cavity myiases (S1 Table), consistent with the anatomical classification of human myiasis reviewed by Francesconi and Lupi [30]. Its development period varies according to ambient temperature. At 37°C, the eggs can hatch rapidly, within 10.5 hours, but it can be prolonged at lower ambient temperatures [14]. This disease may occur concurrently with underlying diseases and the infestations can occur on different body parts with a wide range of clinical symptoms. In our study, we divided the reported human cases into three groups: children aged less than 14-years (Group 0-14), adults aged 15 to 64 (Group 15-64), and the elderly aged 65-years and above (Group 65 and above) (S2 Table).

A total of 173 human cases due to *C. bezziana* worldwide have been recorded with underlying diseases and conditions (Fig 2A, S1 and S3 Table). Among them, open wounds and infections were most commonly recorded. Eighty-seven patients, including 43/92 in Group 65 and above, 39/67 in Group 15-64, and 5/14 in Group 0-14, had open wounds recorded as trauma, ulcer, wound, burns, bed sores, lesion, or orbit postevisceration. Meanwhile, 60 patients, including 28/92 in Group 65 and above, 22/67 in Group 15-64, and 10/14 in Group 0-14, were demonstrated presenting with different infections including serious tropical infectious diseases such as filarial lymphedema, malaria, ankylostomiasis, and leprosy, and also tuberculosis, human immunodeficiency virus (HIV) infection, sepsis, gangrene, pneumonia, chest infection, bronchitis, pleuropneumonia, endophthalmitis, otitis, rhinitis, sinusitis, pansinusitis, herpes simplex stomatitis, dental abscess, chronic pericoronitis, cellulitis, appendicitis, vulvitis and vaginis, hemolytic streptococcal infective endocarditis, herpes zoster opththalmicus, hepatitis B virus infection, perianal codylomata acuminata, aspergillosis, and chromoblastomycosis.

Except for infections and open wounds, the elderly patients in Group 65 above were subject to age-related status as well, with 44 of them being bedridden, wheelchair bound, or debility, 36 even suffering from multiple underlying illnesses, and 18 with tube feeding. The age-related diseases (ARDs) and noninfectious chronic diseases (NCDs) such as dementia (17) and other mental disorder (7), stroke (14), cancer (12), diabetes mellitus (12), and hypertension (10) were commonly recorded. The latter, NCDs, were commonly reported in Group 15-64 as well, including diabetes mellitus (17), cancer (11), mental illness excluding dementia (9) and neurological disorder (3). Multiple underlying illnesses (3) and tracheostomy (3) were also recorded. In addition, miscarriage (1) [31], HIV and hepatitis virus infections complicated with mediolateral episiotomy (1) [32], and postpartum lochia (1) [31], during pregnancy, childbirth and the puerperium were documented among women of childrenbearing age. Otherwise, drug addict (1) [33], chronic seborrhoeic eczema(1) [34] in a German tourist travelling to a Malaysian island in 2012, were also recorded. These data indicated a risk of *C. bezziana* infestation posing to the vulnerable individuals. In Group 0-14, the most commonly reported condition was infections(10), especially ear infections (4/11), and additionally, the susceptible suspects were those children presenting with mental illness excluding dementia (3) and neurological disorder (3), mouth breathing or incompetent lips (1), debility (1), tube feeding(1) and pharyngostomy (1).

**Fig 2.**
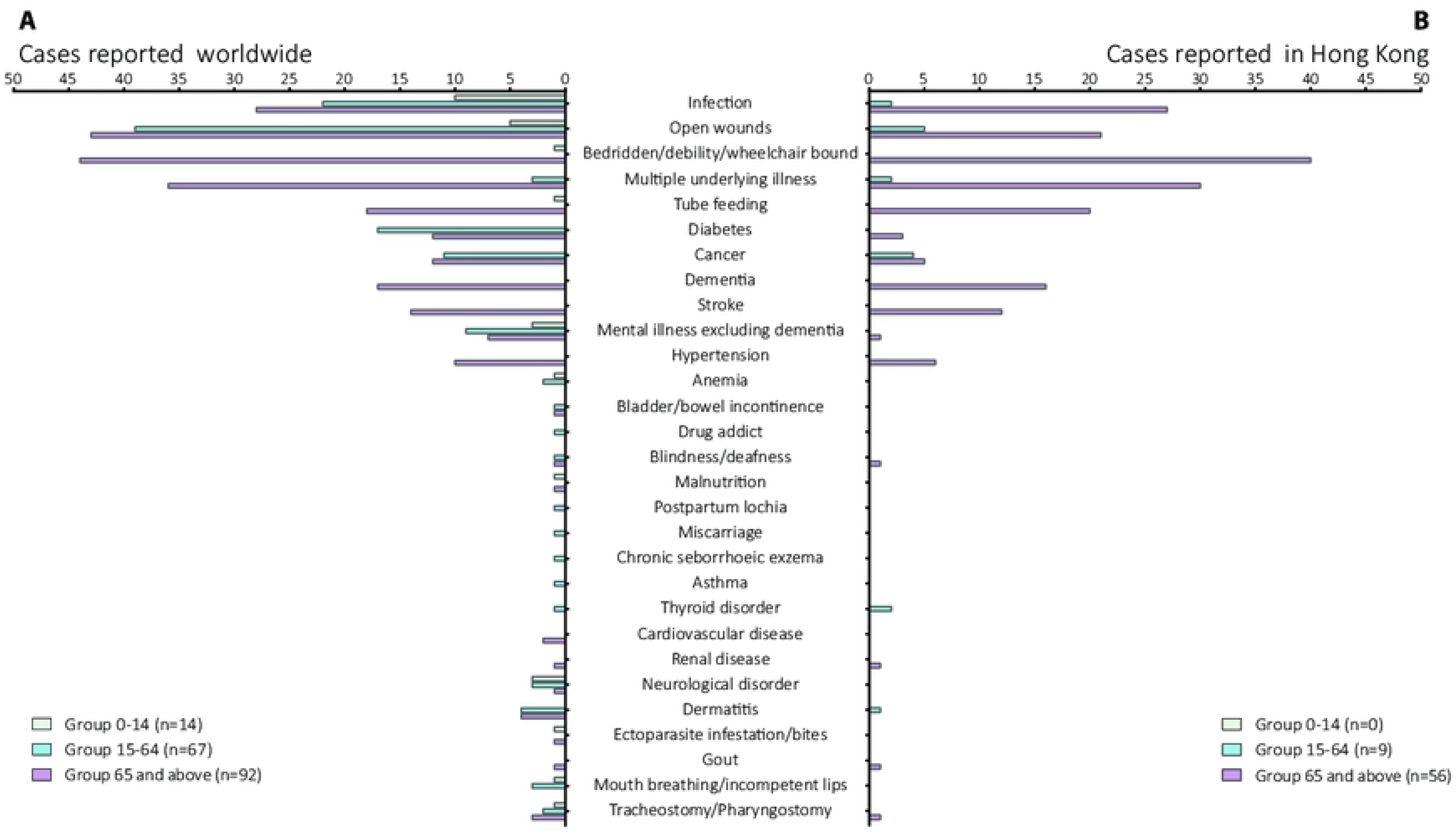
Underlying diseases and conditions recorded in human cases due to *Chrysomya bezziana*. (A) Worldwide. (B) Hong Kong. Data were retrieved from 173 and 65 human cases, worldwide and Hong Kong, respectively. Infection: the infectious diseases caused by different types of pathogens, such as parasites, bacteria, viruses, and fungus; Mental illnesses excluding dementia: described as mental retardation, schizophrenia and related diseases; Neurological disorder: recorded as epilepsy, seizure, neuro-degenerative disorder, cerebral plasy, quadriplegia, kyphoscoliosis, etc.; Ectoparasit infestation/bites: described as pediculosis, and leech bites; Open wounds: included as ulcer, wound, trauma, burns, bed sores, lesion, and orbit postevisceration; Cancer: recorded as cancer, carcinoma, tumor, leukemia, and lymphoma.

Data regarding the site of infestation were summarized from 199 human cases worldwide (Fig 3 and S1 and S4 Table). Among them, 60 and 59 cases of myiasis most commonly occurred in the mouth, and limbs especially the lower limbs. In Group 65 and above, the most common infestation sites were mouth (42/95), limbs (29/95) and eye (11/95). In Group 15-64, limbs (28/72) were the most common site, followed by mouth (12/72), perineal & inguinal regions (9/72), nose (7/72), and torso (7/72) including breast (3), back (1), shoulder (1), buttock (1), and umbilical region (1). While in Group 0-14, the mouth (5/19), ear (4/19), scalp (4/19), and perineal & inguinal regions (3/19) were commonly involved.

More severe complications may occur when myiases emerge in the head and neck regions. In this study, nine cases were recorded the complete destroy of eyes by the aggressive *C. bezziana* larvae in Hong Kong, India, Indonesia, and Iran, five of them suffering from eye cancer [6,22-28], but Sachdev, et al [23] reported a rapid destruction of the globe within two days in a healthy and a non-compromised patient. The destructions of massive tissues and multiple organs involving the eyes, nose, ear and mouth, were also demonstrated and even death came out [6].

Likewise, the loss of function, or even amputation might follow when other parts of the body such as the limbs and/or their appendages are involved [18-21]. In addition, myiases of seven cases occurred in the wounds around tracheostomy or pharyngostomy tubes in a 3-year-old girl and six elderly patients (Figure 2 and S1 Table) [35-40], which put the patients in a risk of a probable airway obstruction or aspiration pneumonitis by aspiration of the screwworms, even the aggressive larvae invasion of the major blood vessels in the neck.

Further analyses of the above data regarding the underlying diseases combined with the infestation sites (S2 Figure and S5 Table), showed that some underlying diseases and conditions more commonly involved some special infestation sites, and vice versa. For example, although infections were described in every site of infestation summarized in our study, the majority of cases involved the main facial organs such as ear (7/8), nose (8/10), and eye (10/15), and even perineal inguinal regions (5/11) were documented. Except for infections, 15 cases of eye infestations were also commonly reported with open wounds (7), cancer (6), multiple underlying illnesses (5), and bedridden, wheelchair bound or debility (5). Four out of five patients involving the face suffered from cancer. Meanwhile, 55 cases occurring in the mouth were most commonly documented with mouth breathing or incompetent lip (4/4), tube feeding (18/19), bedridden, wheelchair bound or debility (33/45), neurological disorder (5/7), dementia (11/17), multiple underlying illnesses (25/39), stroke (7/14), other mental disorder (8/19), infections (16/60), and cancer (7/23). Fifty-seven cases involving the limbs were recorded with dermatitis (7/8), diabetes mellitus (22/29), open wounds (48/87), and infections (12/60). Among them, 22 out of 29 patients with diabetes mellitus most commonly occurred in the low limbs resulting from the diabetic foot (S1 Table) [20,41-43], so it was with the filarial lymphedema in that five out of six patients presented infestation in the lymphedematous limbs (S1 Table) [43-44]. Among 23 underlying cancer patients, the infestation sites commonly occurred in the face (4/5), tracheostomy or pharyngostomy (4/7), eye (6/15), mouth (7/55), and torso (3/9).

**Fig 3.**
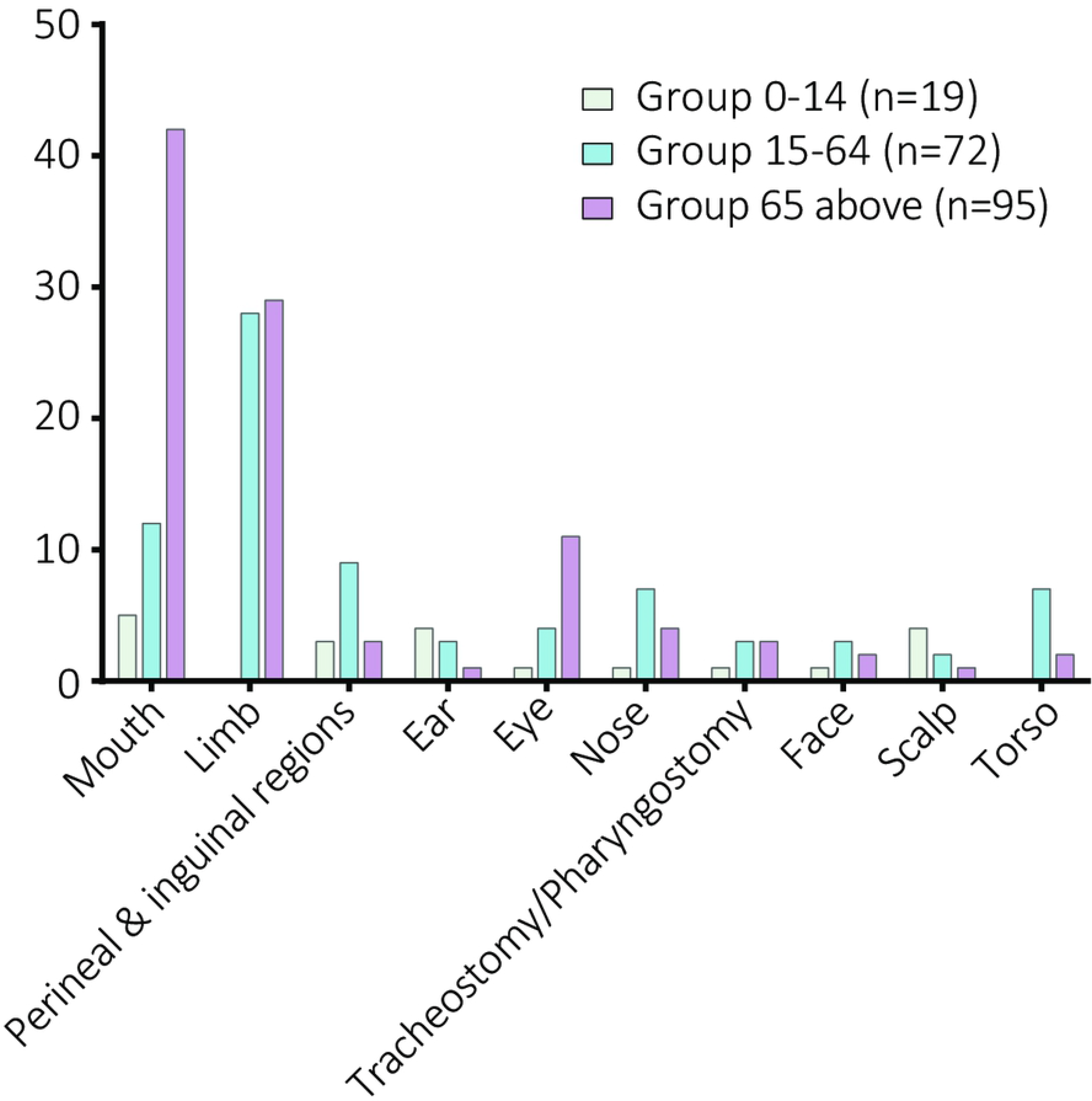
Site of infestation recorded in human cases due to *Chrysomya bezziana* worldwide. Data were summarized from 199 human cases worldwide.

Apart from an ulcer or wound filled with living larvae, the symptoms of *C. bezziana* myiasis were mostly non-specific, ranging from pruritis, pain, to severe tissues and/or bone destruction, recorded in 144 cases (S3 Fig, S1 and S6 Table). The presence of larvae in the lesion and an open wound/ulcer were the most common symptoms among the three groups (120). The other commonly reported symptoms included bleeding (49), ulcer, wound, tunnels or perforation (49), discharge (45), swollen (38), pain (37), fever (30), necrosis (27), severe tissues and /or bone destruction (23), and a foul smell emanating from the wound(21).

The ulcers could get as large as 15×10 cm [45] in size, usually containing larvae as well as serosanguinous fluids. The extensive ulcers may be associated with serious complications such as as loss of limbs and their appendages [19-21,46]. For example, an 89-year-old lady from Hong Kong, who presented to the hospital with an ulcer filled with larvae on her right foot, ended up with below-knee amputation due to the extensive damage caused by *C. bezziana* [46]. In addition, the skin surrounding the lesion was inflamed, with clinical manifestations of swelling, redness, and cellulitis. The discharge was usually purulent, foul smelling and blood stained. Fever was commonly reported in those patients with oral myiasis(16), followed by nasal (7) and limbs (6) myiases. A foul smell, including halitosis, was a common manifestation among the patients with oral cavity myiases.

In our laboratory, we diagnosed a case of wound myiasis caused by *C. bezziana* in an 84-year-old female in Lufeng County, Guangdong Province, China. The lady had a history of hypertension for more than 20 years and had suffered from cerebral thrombosis for 3 years. Four days prior to her admission, she got a scratch with bleeding on her left leg, and at the hospital, the diagnosis of myiasis of the left leg was made. Her main presenting complaints were the sensation of larval movement, skin ulcer and unbearable, intense pain. Her wound quickly progressed from a mere scratch to a 4 cm ×5 cm lesion in five days (Fig 4A). The ulcer became larger and deeper reaching the muscle layer as the insects larvae destroyed the tissues. The patient displayed an increased white blood cell count of 14.14×10^9^/L, a neutrophilic granulocyte (NEU) count of 10.87×10^9^/L, a monocyte count of 0.69×10^9^/L, an eosinophil (EOS) count of 0.61×10^9^/L, a NEU percentage (NEU%) of 76.8%, a C-reactive protein level of 10.7 mg/L and an IgG level of 35.5 g/L. Both *Enterobacter cloacae* and *Stenotrophomonas maltophilia* were isolated from the purulent discharge. This case illustrates the aggressive nature of *C. bezziana* myiasis and the severity of its highly disturbing symptoms.

**Fig 4.**
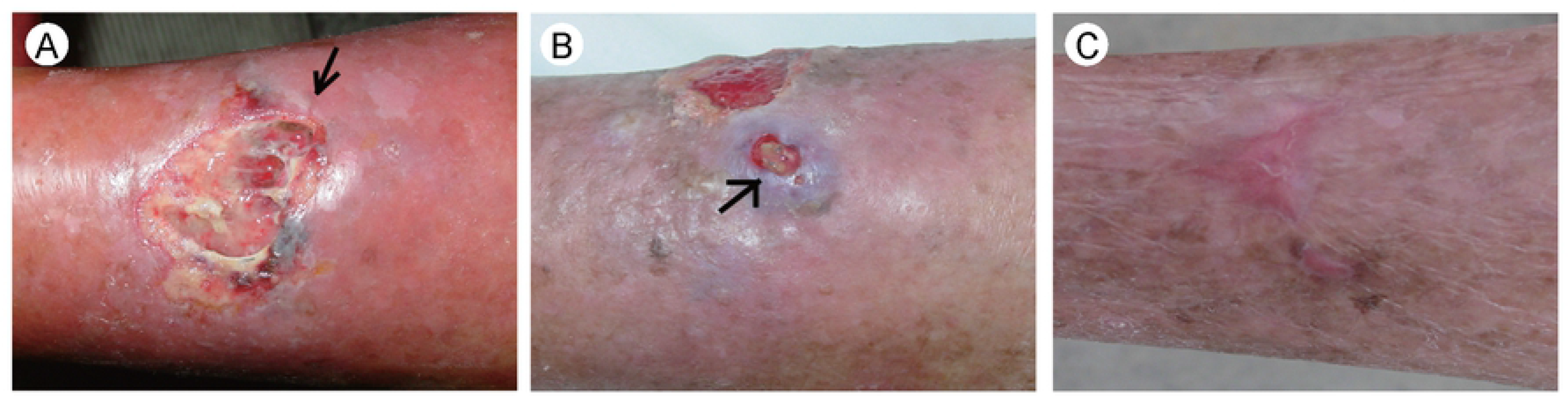
Photographs showing the healing process of the patient’s skin ulcer. (A) The skin ulcer on the patient’s left leg taken on the day of admission, showing a large ulcer measuring 4 cm × 5 cm with undermining up to 8 cm × 8 cm; the black arrow shows tunnels with maggots migrating in. An obvious flare expanded to her whole left limb. (B) The skin ulcer showing a new grayish-blue skin nodule that appeared near the old ulcer, ф1.5 cm; the black arrow shows two larvae in the discharge. (C) The ulcer healed completely.

### Diagnosis

The gold standard for diagnosis of *C. bezziana* myiasis is entomological evidence for species identification. The sampled larvae are killed by immersion in near boiling water (90-100°C) for 30s before being preserved in 70%-95% ethanol (reviewed by Francesconi and Lupi) [47]. The anatomical features of *C. bezziana* can be used for its initial identification: the body shape, body surface, papillae, spiracles (posterior and anterior), dorsal tracheal trunks, and the cephalopharyngeal skeleton (Fig 5A-I) [48-49]. The third stage larvae are 6.1-15.7mm long and 1.1-3.6mm in diameter [2]. The anterior end is armed with paired mouth hooks (mh) which are connected posteriorly with a black cephalopharyngeal skeleton (Fig 5A, B, F and G). The body has prominent bands of thorn-like spines (Fig 5A, B, F-H), the anterior spiracles are occluded with 4-6 papillae (Fig 5G and I), and the pair of posterior spiracles are surrounded each by a peritreme which is incomplete ventrally, each exhibiting three slit-like openings at about 45° to the horizontal (Fig 5C and D). See the diagnostic manual by Spradbery [14] and papers by Sukontason et al [34] and Gan [50] for further details. In the case of the patient diagnosed in our laboratory, dead larvae extracted from the lesion mixed with puss, blood, and mucous discharge can also be used for species identification (Fig 4B and 5E). The dead larvae can be identified by fixing in 70% ethanol, followed by submerging in 10% KOH for 24 hours. This procedure resulted in the effective isolation of the dead larval exoskeletons from the discharge, which assisted species diagnosis (Fig 5E-I).

**Fig 5.**
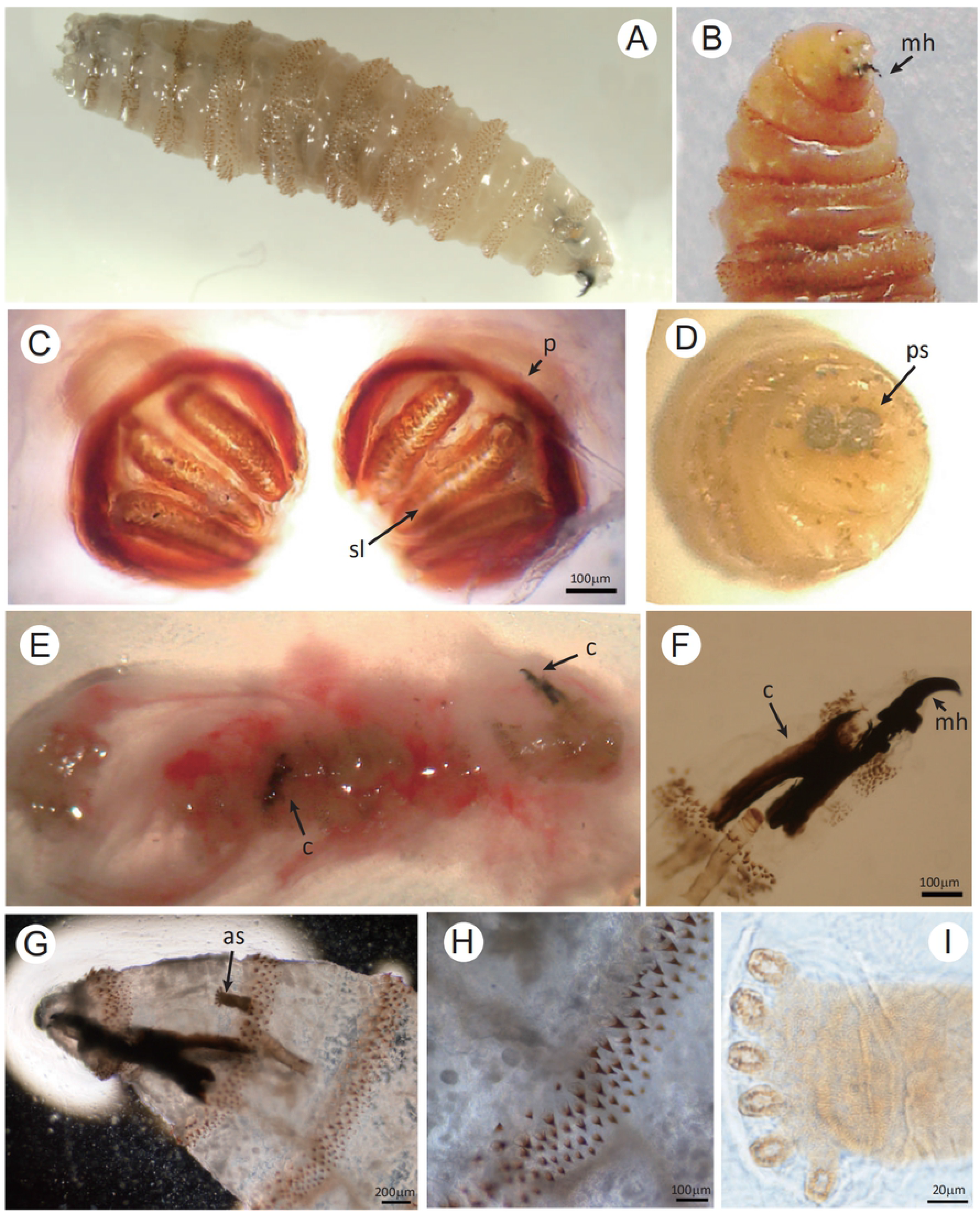
Morphology of the third-instar larva of *Chrysomya bezziana*. (A) The third-instar larva of *C. bezziana* preserved by Food and Environmental Hygiene Department (FEHD), Hong Kong. (B) The larva was sampled from the patient’s skin ulcer after the occlusive treatment using vaseline ointment. Dissecting micrograph of the third-instar larva of *C. bezziana* approximately 14 mm in length, the larva anterior region showing the head region with a strong mouth hook (mh). (C) Light micrograph of the caudal view of the posterior spiracles, which shows that spiracular slits (sl) are slightly convergent and the peritreme (p) is thick and incomplete. (D) Dissecting micrograph of the larval posterior region showing the posterior spiracles (ps). (E-I) Two third-instar larvae of *C. bezziana* were isolated from the discharge extruded from the skin nodule on patient’s left lower limb. (E) Dissecting micrograph of the discharge containing two third-instar larvae with two black cephalopharyngeal skeletons (c). (F, G) Light micrograph of the anterior region of these two third-instar larvae showing the cephalopharyngeal skeleton (c) with strong and robust mouth hooks (mh) which enable the larvae to penetrate deeply into the patient’s muscles, and (G) the anterior spiracle (as) with palmate shape due to six papillae arranged in single row. (H) Light micrograph of enlarged view of the intersegmental spines of the third-instar larva, the single, darkened and tapered tips are recurved toward the body. (I) Light micrograph of enlarged view of the anterior spiracle. (Bar: C, F and H =100 μm; G=200 μm; I =20 μm)

Alternatively, the larvae can be reared to adults followed by morphological identification using the adult taxonomic keys [14,51]. Meanwhile, the real-time Polymerase Chain Reaction (PCR) techniques is widely used not only for adult fly surveillance identifications, but also to confirm the identity of *C. bezziana* [52-53]. Computer tomography (CT) or magnetic resonance imaging (MRI) have been mainly utilized to locate the larvae and to delineate the extent of the erosion, especially when the disease involves sensitive body parts such as the mouth, nasopharynx, eye, ear, etc. [25,54-55].

### Treatment

The most important and effective treatment is the removal of all living larvae quickly and thoroughly [47]. Alhady et al reported that a 9-year-old boy presenting severe ophthalmomyiasis from *C. bezziana* resulted from his primary aural myiasis, warning of an early and thorough larvae removal [56]. Treatment options depend on the different sites of infestation and different degrees of damage (S1 and S7 Table). Firstly, manual removal of all visible larvae can be carried out using tweezers or forceps followed by debridement to remove the necrotic tissues. This method can be used to treat myiasis affecting the majority of infested body sites (S7 Table). For a further option, paralyzing agents for local application, such as chloroform, ether and 1% lidocaine, are used to facilitate larval removal [6,3,57-60]. Secondly, surgical removal that involves extensive wound exploration under anesthesia may be required depending on the degree of tissue damage. The Infestations in the mouth (18), limbs (12), eyes (10), ear (3), and tracheostomy/pharyngostomy wounds (3) were treated by surgery (S7 Table). Fourteen of them were recorded with serious necrosis, severe tissues and/or bone destruction, nine patients received the eye exenteration [6,23-28,61], and four underwent the limb amputation [18-21](S1 Table). Thirdly, the use of suffocating agents such as turpentine oil, butter, mineral oil, vaseline, liquid paraffin, petroleum jelly, bee wax, resins, chewing gum, pork fat and polymyxin B sulfate ointment (reviewed by Nene et al [62]) are convenient and effective for patients, especially at the primary care level. These agents force the larvae out by blocking air entry via the spiracles so that they can easily be picked up by forceps or tweezers [62]. This method proved to be effective where the larvae were successfully removed from the most infested sites, even complex deep structures, including limb (13), mouth (11), eye (6), nose (4), perineal & inguinal regions (4), face (3), ear (2), scalp (2), and even the tracheostomy or pharyngostomy wounds (S1 and S7 Table). The turpentine oil was commonly used as the effective suffocating agent. The cotton buds or gauze impregnated with turpentine oil were usefully applied in the myiases treatment involving limbs, mouth, eye, ear, nose, facial tumor base [33,62-65], and perineal & inguinal regions [58,66]. If the oclusive measure was applied to the infested sites around the tracheostomy and pharyngostomy tubes, gave great caution for the high risk of chemical pneumonitis [67]. In one treat, the turpentine liniment was used in the scalp [68], In another, Bayer [69] reported that plugs soaked in 10% dextrose solution were successfully used to remove larvae from ear and nasal myiases due to *C. bezziana*, as the larvae seemed to have a liking for the dextrose solution, and consequently invade the plugs and stick to it, which faciliated larvae removal. Therefore, it is recommended to be applied on other infested parts as well. Finally, the use of entomological insecticides such as ivermectin has been recorded in India [64-65,70-74].

Apart from complete removal of all living larvae, antibiotic therapy, nutrition support, and maintenance therapies against any underlying diseases should be prioritized for those patients in need. The antibiotics were prescribed to treat and prevent bacterial secondary infections. Nutritional support should be considered for those patients with poor general condition and those with serious physical damage. Early and timely treatment is essential to help wound healing and stop the rapid and destructive process of *C. bezziana* myiasis. However, some patients have a bad prognosis so that severe damage to the tissues cannot be restored in the short-term, especially those suffered with serious underlying diseases and myiases affecting sites such as the eyes, ears, mouths, noses, throats and other organs or soft tissues [6,23].

In the case of the patient diagnosed in our laboratory, the treatment involved multiple therapies of larvae removal by the vaseline ointment occlusion method, wound debridement, antibiotic and anti-hypertensive treatment as well as nutritional support. The vaseline ointment was promptly smeared over the ulcer and applied at more than five mm thick for 24 hours. After only half an hour, the patient’s pain was alleviated and her condition relieved significantly. Eight larvae crawled out of the ulcers one after another in 10 hours. We removed the visible larvae carefully by using tweezers. After treatment, the nodule beside the ulcer (Fig 4B) was destroyed, and two dead larvae were discharged (Fig 5E). The patient was managed promptly with experienced doctors in our hospital and the myiasis ulcers with secondary bacterial infection were completely healed after three weeks of antibiotic treatment. Complete wound healing took about three months (Fig 4C). In the case of the frail 84-year-old lady, her invasive injuries could have been curtailed quickly and her suffering would have been reduced, if prompt diagnosis and proper treatment was given at the rural clinic.

### Outcomes

In this study, 171 patients had the documented outcomes which varying with their health status and the infested sites (S1 and S8 Table). The majority (148) of them had positive outcomes after effective treatment, but 23 died (S8 Table). Even among the majority (148), four patients had received limbs amputation [18-21], and seven eye exenteration [23-28] due to the aggressive tissues and/or bone destruction of *C. bezziana* larvae. Among the death cases (23), 22 occurred in Group 65 and above, only one in Group 15-64. Twenty out of 23 were recorded with multiple underlying diseases, including the patient in Group 15-64, who was a 39-year-old Indian woman suffering from multiple underlying illnesses including malaria, ankylostomiasis, etc., and her death was attributed to the worsened multiple underlying diseases and the extreme exhaustion with rapid and heavy destroy in the main facial organs caused by thousands of screwworms with several generations (S1 Table) [6]. The infested sites in the death patients were commonly reported in the mouth (20/23), followed by eyes (2/23), nose (2/23), face (1), and torso (bed sores, 1) (S8 Table). The most common cause of death was recorded as pneumonia (10/23), followed by myocardial infarction (3/23) and sepsis (3/23). Therefore, clinicians should bear in mind that prompt and proper management of both myiasis and multiple underlying diseases, especially for those elderly patients, is key to improved outcomes.

### Epidemiology

#### Human myiasis reports worldwide

From 1909 to 2019, 16 countries worldwide have recorded at least 291 human cases of *C. bezziana* myiases (Table 1, S1 and Fig 6A). According to the published literature, Asia has reported by far the highest rate of cases 94.5% (275/291), most cases were distributed in China 36% (99/275) (85 cased recorded in Hong Kong, 14 in mainland China) (Fig 6B), and India 36% (99/275), followed by Sri Lanka 15.6% (43/275), and Iran 4.7% (13/275), among them, 111 out of 275 reported in the recent ten years in Asia. Meanwhile, Oceania and Africa only reported 3.1% (9/291) and 2.4% (7/291) human cases, respectively, and all 7 human cases in Africa and 6 in Oceania were recorded before 1968. Although there have been reports of *C. bezziana* associated with human myiasis in Algeria, northern Africa [9], Turkey [10], Spain [11] and Mexico [12] but in all these cases the identity of the myiasis larvae might be incorrect; the most likely species being *Wohlfahrtia magnifica* in Europe and Africa and the New World screw-worm fly, *Cochliomyia hominivorax*, in Mexico.

**Fig 6.**
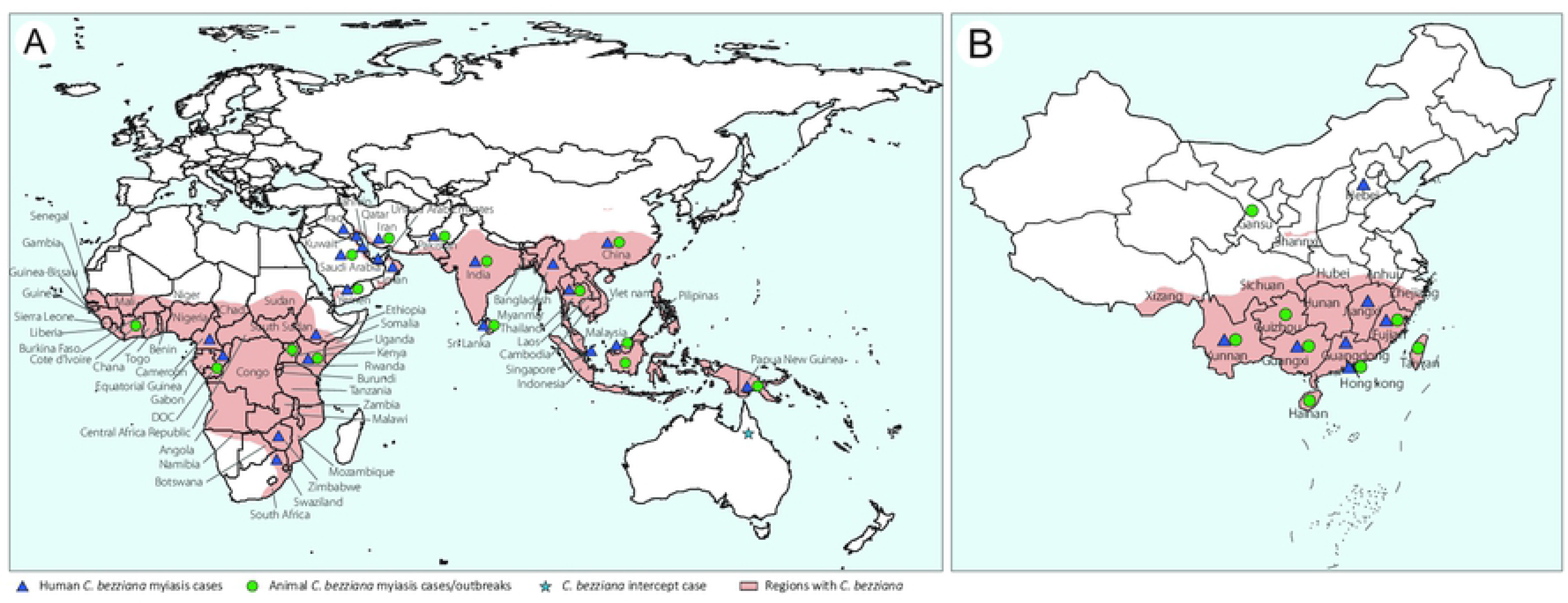
Geographical distribution of species *Chrysomya bezziana* and myiasis caused by *C. bezziana*. (A) Worldwide; (B) China. The maps were conducted using AI.

**Table 1.** Number of human cases reported in each continent in different time periods.

| Country/Region | Continent | -1968 | 1969-1978 | 1979-1988 | 1989-1998 | 1999-2008 | 2009-2019 | Total |
| --- | --- | --- | --- | --- | --- | --- | --- | --- |
| China | East Asia | 3 | 4 | 4 | 0 | 42 | 46 | 99 |
| India | South Asia | 60 | 1 | 0 | 7 | 7 | 24 | 99 |
| Sri Lanka | South Asia | 2 | 0 | 0 | 14 | 0 | 27 | 43 |
| Iran | Middle East | 0 | 1 | 0 | 1 | 3 | 8 | 13 |
| PNG | Oceania | 6 | 0 | 3 | 0 | 0 | 0 | 9 |
| Yemen | Middle East | 0 | 0 | 0 | 0 | 8 | 0 | 8 |
| Malaysia | Southeast Asia | 0 | 0 | 1 | 0 | 3 | 2 | 6 |
| Uganda | Africa | 3 | 0 | 0 | 0 | 0 | 0 | 3 |
| DRC | Africa | 2 | 0 | 0 | 0 | 0 | 0 | 2 |
| Indonesia | Southeast Asia | 0 | 0 | 0 | 0 | 1 | 1 | 2 |
| Saudi Arabia | Middle East | 0 | 0 | 2 | 0 | 0 | 0 | 2 |
| Cote d'Ivoire | Middle East | 1 | 0 | 0 | 0 | 0 | 0 | 1 |
| Kenya | Africa | 1 | 0 | 0 | 0 | 0 | 0 | 1 |
| Laos | Southeast Asia | 0 | 0 | 0 | 0 | 0 | 1 | 1 |
| Pakistan | South Asia | 0 | 0 | 0 | 0 | 0 | 1 | 1 |
| Thailand | Southeast Asia | 0 | 0 | 0 | 0 | 0 | 1 | 1 |
| Total |  | 78 | 6 | 10 | 22 | 64 | 111 | 291 |

PNG=Papua New Guinea, DRC=Democratic Republic of Congo

In the present study, 190 patients had recorded with age (S2 Table), 193 with gender (S9 Table), and 165 with socioeconomic status (S10 Table). Worldwide, *C. bezziana* myiasis commonly affects old people, as 96 out 190 human cased recorded in Group 65 and above, 73 in Group 15-64, and 21 in Group 0-14 (S2 Table). While 154 out of the 165 (93.3%) patients were of low socioeconomic status or living in the elderly home, etc (S10 Table). But no obvious differences were found in gender analyses both in the world and Hong Kong (S9 Table).

In Hong Kong, the first case of human myiasis due to *C. bezziana* was reported in 2002 [61]. Since then, the government of Hong Kong, through the Center of Health Protection (CHP) has been reporting the monthly occurrence of cases consistently through the periodical known as *Communicable Diseases Watch (CDW)*. Although *C. bezziana* myiasis is not a notifiable disease in Hong Kong, this publication provides a viable opportunity for a better understanding of the epidemiology status in a city. *C. bezziana* myiasis-associated data are available in Hong Kong from both government documents (CDWs) and the published literature, where at least 85 human cases have been recorded since 2002. We believe there may well have been *C. bezziana* myiasis in Hong Kong before it was first reported in 2002. From the data available, the cases appear to present a predilection for Group 65 and above, in which 83.8% (56/68) of cases occurred, with a mean age of 82.6 years (S2 Table). The most common underlying diseases among 56 elderly patients in Group 65 and above in Hong Kong were infections, ARDs and NCDs, 30 suffering with multiple underlying illnesses, 40 under bedridden or wheel chair bound or debility status, 27 with infections, 16 with dementia, 12 with stroke, 20 assisted with tube feeding, meanwhile 21 patients having open wound (Fig 2B, S1 and S11 Table). Moreover, Hong Kong is a developed city and has an aging society. In 1996 and 2016, the population of aged (the percentage older than 65 and above) in Hong Kong reached 10.1% and 16%, respectively, and it is expected to increase to 34% by 2066 [75]. Therefore, the urgency for preventing myiasis due to *C. bezziana* for the elderly people, especially aged 65 and above suffering from infections, ARDs or NCDs, being debilitated and living in the elderly home, cannot be underestimated.

#### Animal myiasis worldwide

Since 1909 [4], animal cases and outbreaks due to *C. bezziana* have been recorded in 24 countries worldwide, 16 of them from Asia, including China, Saudi Arabia, Iran, India, Iraq, Oman, Yemen, Pakistan, Kuwait, Singapore, Sri Lanka, Myanmar, Indonesia, United Arab Emirates (UAE), Bahrain and Malaysia (S12 Table and Fig 6A), 7 from Africa, including Zimbabwe, Cameroon, Chad, Kenya South Africa, Ethiopia, and Congo (DRC) [2], and the last from Papua New Guinea, Oceania, which was sourced from investigations by Spradbery et al. (2019) [76]. At least 19 species of domestic and wild animals have been reported to be affected by *C.bezziana* including cattle, sheep, goats, dogs, horses, camels, buffaloes, donkeys, pigs, Sambar and Rusa deer, water buffalo, cats, birds, wallabies, cus cus, rabbits, tree kangaroos, elephant, and broilers (S12 Table). These animals, especially the domesticated species, may be a high risk in transmitting the disease to humans. In China, outbreaks of the disease among animals have been recorded in 8 provinces/province-level regions including Hainan, Fujian, Guangxi, Yunan, Guizhou, and Gansu, as well as Hong Kong and Taiwan (Fig 6B and S12 Table). From 1987 to 1991, 513 animal cases occurred in Guizhou Province. In Fujian Province, there were 20 and 85 cattle cases, and 3 pig cases reported in 2001, 2007 and 2011, respectively. Meanwhile, in recent ten years, animal myiasis outbreaks due to *C. bezziana* were reported in Indonesia in 2006-2009, 2009-2012 and 2014, Kuwait in 2008, Yemen in 2008-2009, Pakistan in 2012-2015, Iraq in 2013-2014, Sri Lanka in 2014, India in 2014 and 2018, Saudi Arabia in 2015-2016, Malaysia and Singapore in 2017 (S12 Table).

### Species distribution

The distribution of *C. bezziana* has been described by Animal Health Australia (AHA, 2017) and CABI (https://www.cabi.org/ISC/), an approximation for the global distribution was plotted involving 65 countries in Asia, the Indian subcontinent, the Middle East, PNG, tropical and subtropical Africa, especially in sub-Saharan Africa (Fig 6A) [2]. In the present study, the occurrence of *C. bezziana* has been summarized in at least 44 countries worldwide in the published literature (S13 Table, Fig 6A). Meanwhile, *C. bezziana* has been intercepted by quarantine in Australia [77]. However, there is a possibility of permanent colonization of new geographical areas by this species after accidental introduction, either through an infested host, or aircraft or returning livestock vessel [78]. For instance, some countries in the Middle East, such as Bahrain, Kuwait, Iraq and Yemen have been colonized by *C. bezziana* after incidental introductions [79]. In addition, the spread of human myiasis cases may be associated with aircraft travel [79].

In this study, the recorded data indicated that *C. bezziana* was found in 17 provinces/province-level regions in China, including Hainan, Fujian, Guangdong, Guangxi, Yunan, Guizhou, Sichuan, Qinghai, Xizang, Hunan, Hubei, Jiangxi, Gansu, Shannxi, Hebei, as well as Hong Kong and Taiwan(Fig 6B and S13 Table). Its distribution pattern in China is mainly the tropical-subtropical areas of the Oriental region. But animal outbreaks and human cases were recorded in only 8 and 6 provinces/province-level regions such as Hong Kong, respectively. Although 85 human cases have been reported in Hong Kong, only 14 human cases have been reported in mainland China, including the provinces of Fujian [80], Guangdong, Guangxi [57,81], Yunnan [3,81], Jiangxi [82], and Hebei [83] (Fig 6B and S1 Table).

Therefore, both in China and the world at large, it is possible that *C. bezziana* distribution could be far greater than currently reported. Sutherst et al [78] predicted that some parts of the Americas and Australia could provide favorable conditions for colonization by *C. bezziana* once introduced. The potential for incursions and subsequent spread of *C. bezziana* poses a risk to global public health. Furthermore, systematic surveillance and treatment studies are required and these would be expected to provide for the deployment of better *C. bezziana* prevention, control and treatment strategies worldwide.

### Prevention and Control

The present study has suggested that *C. bezziana* myiasis commonly affects patients having underlying diseases, especially the elder, individuals of low socioeconomic status, and those with poor personal hygiene. This review has shown that the disease in humans most commonly occurs in the mouth, followed by the limbs. In animals, cases are mainly recorded in domesticated species, including pets. Therefore, the prevention and control of *C. bezziana* myiasis requires an integrated approach that includes personal protection, environmental improvement, good animal husbandry practices, proper keeping of pet animals such as dogs and legislation including making the condition a notifiable disease.

We highlights the urgent need of health education and supervision of personal protective measures to those patients, their families, and the health staff, especially in the elderly homes or the primary health care centers, in preventing *C. bezziana* infestation. These measures can be applied in order to block female flies from laying eggs on humans, especially on the elder, the bedridden or debiliated, those who suffered from infections, ARDs, or NCDs, patients with different types of open wounds including trauma, burns, ulcers, diabetic foot, bed sores, orbit postevisceration, lymphedematous limbs, wounds around tracheostomy or pharyngostomy tubes, and etc. Infestations in children, especially those affecting their mouth, ears, scalp, and perineal & inguinal regions must be diagnosed and treated promptly to control the risk of contracting myiases in this vulnerable age group. In addition, those with miscarriage [81], mediolateral episiotomy [66], and postpartum lochia [81] in women of childrenbearing age, drug addict [84], tourist travelling to the epidemic area [85] also need a particular concern. It is essential to maintain good personal hygiene, such as skin and oral care, particularly for those who do not have the ability to take care of themselves. Since blood, wound exudates and smell/odour are known to attract the gravid female flies to deposit their eggs on a host [30], all types of wounds need to be kept clean and thoroughly dressed, in particular, for the patients leaving hospitals with cancerous lesion, tubes of feeding, tracheostomy or pharyngostomy, and etc.

Physical barriers can be employed to protect individuals at risk. Such measures include using bed nets [30], and stationing screens in all possible points of entry such as air vents, windows and doors [87]. The use of anti-fly curtains, air curtains, and insect electrocutors are also recommended [87]. In addition, the appropriate use of insecticides [88] can also be used to reduce the risk of *C. bezziana* myiasis.

Domesticated animals including pet dogs may act as fly attractants if they have not been properly kept and as a result, those individuals working in animal husbandry, as well as pet owners may be at risk of contracting *C. bezziana* myiasis. Therefore, deliberate efforts should be made to ensure that domestic animals including pets are regularly and carefully inspected by owners and veterinary personnel so that *C. bezziana* can be intercepted promptly.

Respective governments can play a key role at the national level in the fight against *C. bezziana*. For instance, the government can impose quarantine restrictions on *C. bezziana* cases from endemic areas. To achieve this, however, functional surveillance systems must be in place. This measure has been illustrated by the Australia Government which, through functional surveillance systems and quarantine restrictions, has prevented *C. bezziana* incursions into Australia [89]. Governments can embark on fly eradication programs by using strategies that have proven successful in eradicating the related New World screw-worm fly, the most effective being the sterile insect technique (SIT) [90]. In addition, health personnel should ensure that health care workers, farmers, and the general public are aware of *C. bezziana* in order to enable prompt diagnosis and treatment.

### Limitation

Our study has such limitations as (1) except Animal Health Australia and Hong Kong, data only available from the published literatures, could not from the authoritative government appraisal agency, (2) paucity of studies on larger and systematic surveillance of *C. bezziana*.

## Conclusion

This is a first in-depth study on human myiasis due to *C. bezziana*. Our research provides an opportunity for clinicians to gain a better understanding of this disease from its etiology to its prevention and control. We summarized 291 human cases worldwide and found that theses cases commonly occurred in poor hygiene, low socio-conomic status, old age, and underlying diseases including infections, ARDs, and NCDs. *C. bezziana* myiasis is a devastating and rapidly-progressing condition, posing a risk to public health. Pitifully, however, this disease appears to have been under-recognized as a serious medical or veterinary condition, for human and animal cases have only been reported in 16 and 24 countries respectively though it is recorded to be present in 44 countries worldwide. *C. bezziana*, an obligate parasite, is unable complete its life cycle without parasitizing a warm-blooded host. Cryptic cases do occur and may well be under-recognized or unreported. Through this comprehensive study on human *C. bezziana* myiasis, we call for vigilance from governments, health authorities, health staff, veterinary personnel, the elderly homes, and also the general public.

## Supporting information

S1 Fig. The PRISMA flowchart.

(PDF)

S2 Fig. Heat map of the underlying diseases combined with the infestation sites.

(PDF)

S3 Fig. Clinical signs and symptoms of human cases due to *Chrysomya bezziana* myiasis.

(PDF)

S4 Fig. Preferred Reporting Items for Systematic Reviews and Meta-analysis (PRISMA) checklist.

(PDF)

S1 Table. Characteristics of human cases due to *Chrysomya bezziana* myiasis recorded worldwide.

(PDF)

S2 Table. The patients with *Chrysomya bezziana* myiasis were grouped by age.

(PDF)

S3 Table. Underlying diseases and conditions in patients with myiasis due to *Chrysomya bezziana* worldwide.

(PDF)

S4 Table. Sites of infestation recorded in human cases due to *Chrysomya bezziana* worldwide.

(PDF)

S5 Table. Summary of underlying diseases and conditions combined with sites of infestation in patients with myiasis due to *Chrysomya bezziana* worldwide.

(PDF)

S6 Table. Clinical signs summarized from human cases due to *Chrysomya bezziana* worldwide.

(PDF)

S7 Table. Summary of main therapies recorded in human cases with *Chrysomya bezziana* myiasis worldwide.

(PDF)

S8 Table. Health outcomes of human cases due to *Chrysomya bezziana* worldwide.

(PDF)

S9 Table. A gender analysis in patients with *Chrysomya bezziana* myiasis.

(PDF)

S10 Table. Analysis of the socioeconomic status of patients with *Chrysomya bezziana* myiasis.

(PDF)

S11 Table. Underlying diseases and conditions in patients with myiasis due to *Chrysomya bezziana* in Hong Kong.

(PDF)

S12 Table. Characteristics of animal cases or outbreaks due to *Chrysomya bezziana* myiasis recorded worldwide.

(PDF)

S13 Table. Distribution of *Chrysomya bezziana* recorded in the world.

(PDF)

## Acknowledgement

We give special thanks to Prof. Geqiu Liang (Sun Yat-sen University, China) and Prof. Mengyu Zhang (Hebei Medical University, China) for assistance with species identification, Prof. Qing Chen (Department of Epidemiology, School of Public Health, Southern Medical University, China) for her guidance in systematic review methodology. This study was supported by the National Key Research and Development Program of China (2016YFC1200500), the Guangzhou Synergy Innovation Key Program for Health (201803040006 and 201508020263), and the Guangzhou International Science and Technology Cooperation Program (2012J5100026). The funders had no role in study design, data collection and analysis, decision to publish, or preparation of the manuscript.

## Ethics Statement

The patient in this manuscript has given written informed consent (as outlined in the PLOS consent form) to publication of their case details.

## Author Contribution

Conception and design: XHZ and XYZ; photographing, drawing and chart making: XG, XYZ, DX, YHX, GYD, DMK and TX; clinical management of the patient and specimen collection: XYZ, STZ, YMY and LL; specimen preparation: XHZ, QZM and RJM; species identification: FLJ, XHZ, MCY and PS; data collection and analyses: XHZ, XYZ, PS, MCY, DMK, XG, STZ, DX, YMY, JL, WXW, LXX, STL, YQF, SGH, YXH, and XQZ; results explanation: XHZ, XYZ, PS, MCY and FLJ; drafting: XHZ, XYZ, DMK, STZ, XG, PS, MCY, DX, YMY, JL, WXW, LXX, STL and SPK; editing: XHZ, DMK, XG, RKW, PS, MCY, XYZ and STZ. All authors have read and approved the final version of the manuscript.

## References

1. Scruggs CG. The Peaceful Atom and the Deadly Fly. Austin, Texas: The Pemberton Press; 1975.

2. AHA. Old World Screw-Worm Fly: A Diagnostic Manual, 3rd Ed. Canberra: Animal Health Australia; 2017.

3. Fan ZD. Fauna Sinica Insecta Vol. 6 Diptera: Calliphoridae. Beijing: Science Press; 1997: 55–159.

4. Rovere J. Etude de larvae cuticoles appartenant au genre Chrysomyia, observées au Congo Belge. Bulletin Agricole du Congo Belge. 1910;1: 26–35.

5. Centre for Agriculture and Biosciences International (CABI). Chrysomya bezziana (Old World screw-worm). Avaliable from https://www.cabi.org/isc/datasheet/88417.

6. Patterson RL. An Indian screw-worm. Ind Med Gaz. 1909;44(10): 374–376.

7. Anon. A national review of Australia’s longer term screw fly (SWF) preparedness strategy. Canberra: D.P.I.E; 1990: 32.

8. Spradbery JP. Screw-worm fly: a tale of two species. Agricultural Zoology Review. 1994;6: 1–62.

9. Abed-Benamara M, Achir I, Rodhain F, Perez-Eid C. First Algerian case of human otomyiasis from *Chrysomya bezziana*. Bull Soc Pathol Exot. 1997;90(3): 172–175.

10. Casa I, Satar A, Ünlü K, Sakalar YB, Ari S. External ophthalmonmyiasis infestation. Jpn J Ophthalmol. 2006;50: 176–190. doi: 10.1007/s10384-005-0282-0.

11. Aguado Lobo M, Hernández-Núnez A, Isabel García-Arata M, Borbujo J. Autochthonous cutaneous myiasis due to *Chrysomya bezziana*. Actas dermo-sifiliograficas. 2014;105(5): 522–524. doi: 10.1016/j.ad.2013.07.005.

12. Romero-Cabello R, Calderón-Romero L, Sánchez-Vega JT, Tay J, Romero-Feregrino R. Cutaneous myiasis caused by *Chrysomya bezziana* larvae, Mexico. Emerg Infect Dis. 2010;16(12): 2014–2015. doi: 10.3201/eid1612.100938.

13. Zumpt F. Myiasis in man and animals in the Old World. London: Butterworths; 1965.

14. Spradbery JP. A manual for the diagnosis of screw-worm fly. Canberra: Department of Agriculture Fisheries and Forestry; 2002.

15. Yan RL, Wong Y. Studies on *Chrysomya bezziana* Villeneuve and the myiasis it produced. Wuyi Science Journal. 1983;3: 157–163.

16. Spradbery JP, Ford RJ, Tozer RS. Diel larval exodus in the screw-worm fly, *Chrysomya bezziana*(Villeneuve). J Aust Entomol Soc. 1983;22: 261–262.

17. Chemonges-Nielsen S. *Chrysomya bezziana* in pet dogs in Hong Kong: a potential threat to Australia. Aust Vet J. 2003;81(4): 202–205. doi: 10.1111/j.1751-0813.2003.tb11471.x.

18. Center for Health Protection. A case of human myiasis. Communicable disease watch. 2012;9: 58.

19. Chan JC, Lee JS, Dai DL, Woo J. Unusual cases of human myiasis due to Old World screwworm fly acquired indoors in Hong Kong. T Roy Soc Trop Med H. 2005;99(12): 914–918. doi: 10.1016/j.trstmh.2005.06.020.

20. Amandeep S, Devinder S. Wound myiasis due to *Chrysomya bezziana* (Diptera: Calliphoridae) in patients of diabetic foot. Journal of Entomological Research. 2006;30: 367–369.

21. Rahoma AH, Latif B. Human foot myiasis in Malaysia with a review of the literature. J Trop Med Parasitol. 2010;33: 41–43.

22. Yeung JC, Chung CF, Lai JS. Orbital myiasis complicating squamous cell carcinoma of eyelid. Hong Kong Med J. 2010;16(1): 63–65.

23. Sachdev MS, Kumar H, Roop, Jain AK, Arora R, Dada VK. Destructive ocular myiasis in a noncompromised host. Indian J Ophthalmol. 1990;38: 184–186.

24. Berenji F, Hosseini-Farash BR, Marvi-Moghadam N. A Case of Secondary Ophthalmomyiasis Caused by *Chrysomya bezziana* (Diptera: Calliphoridae). J Arthropod Borne Dis. 2014;9(1): 125–130.

25. Khataminia G, Aghajanzadeh R, Vazirianzadeh B, Rahdar M. Orbital Myiasis. Journal of Ophthalmic & Vision Research. 2011;6: 199–203.

26. Radmanesh M, Khataminia G, Eliasi P, Korai MK, Ebrahimi A. *Chrysomyia bezziana* infested basal cell carcinoma destroying the eye. Int J Dermatol. 2000;39: 455–462.

27. Yaghoobi R, Bagherani N. *Chrysomya bezziana* infestation in a neglected squamous cell carcinoma on the face. Indian journal of dermatology, venereology and leprology. 2009;75: 81–82.

28. Kersten R, Shoukrey N, Tabbara K. Orbital Myiasis. Ophthalmology. 1986;93(9): 1228–1232. doi: 10.1016/S0161-6420(86)33592-9.

29. Humphrey JD, Spradbery JP, Tozer RS. *Chrysomya bezziana*: Pathology of Old World screw-worm fly infestations in cattle. Exp Parasitol. 1980;49(3): 381–397. doi: 10.1016/0014-4894(80)90073-9.

30. Francesconi F, Lupi O. Myiasis. Clin Microbiol Rev. 2012;25(1): 79–105. doi: 10.1128/CMR.00010-11.

31. Center for Health Protection. A case of human *Chrysomya bezziana* infestation confirmed. Feature Topic, March, 2006. Avaiable from: https://www.chp.gov.hk/en/features/6016.html#(accessed 2018).

32. Spradbery JP, Kirk J. Incidence of Old World screw-worm fly in the United Arab Emirates. The Vet Rec. 1992;130(2): 33. doi: 10.1136/vr.130.2.33.

33. Bhola N, Jadhav A, Borle R, Adwani N, Khemka G, Jadhav P. Primary oral myiasis: a case report. Case Reports in Dentistry. 2012;2012: 734234. doi: 10.1155/2012/734234.

34. Sukontason KL, Piangjai S, Boonsriwong, Bunchu N, Ngern-klun R, Vogtsberger RC, Sukontason K. Observations of the third instar larva and puparium of *Chrysomya bezziana* (Diptera: Calliphoridae). Parasitol Res. 2006;99: 699. doi: 10.1007/s00436-006-0233-9.

35. Center for Health Protection. Five sporadic cases of human myiasis. Communicable disease watch. 2010;7: 101–102.

36. Center for Health Protection. A case of human myiasis. Communicable disease watch. 2012;9: 54.

37. Hemanth V, Kumar CS, Manikandan D, Musarrat F, Preetham AP, Paulraj MG. An unusual cause of late tracheostomy bleed. Sci Res. 2013;2(4): 260–262.

38. Gopalakrishnan S, Srinivasan R, Saxena SK, Shanmugapriya J. Myiasis in different types of carcinoma cases in southern India. Indian journal of medical microbiology. 2008;26: 189–192.

39. Prasanna Kumar S, Ravikumar A, Somu L, Prabhu P, Subbaraj R. Tracheostomal myiasis: a case report and review of the literature. Case reports in otolaryngology. 2011;2011: 303510.

40. Soleimani-Ahmadi M, Vatandoost H, Hanafi-Bojd AA, Poorahmad-Garbandi F, Zare M, Hosseini S. First report of Pharyngostomy wound myiasis caused by *Chrysomya bezziana* (Diptera: Calliphoridae) in Iran. Iran J Arthropod Borne Dis. 2013;7(2): 194–198.

41. Faramarzi A, Rasekhi A, Kalantari M, Hatam G. *Chrysomya bezziana* as a causative agent of human myiasis in Fars Province, Southern Iran. Iran J Arthropod Borne Dis. 2009;3(1): 60–63.

42. Kumarasinghe SP, Karunaweera ND, Ihalamulla RL. A study of cutaneous myiasis in Sri Lanka. Int J Dermatol. 2000;39: 689–694.

43. Bambaradeniya Y, Karunaratne W, Rakinawasam S, Tomberlin J, Goonerathne I, Kotakadeniya R. Myiasis incidences reported in and around central province of Sri Lanka. Int J Dermatol, 2019:58(3): 336–342. doi: 10.1111/ijd.14291.

44. Radhakrishnan R, Srinivasan R, Krishnamoorthy K, Sabesan S, Pani SP. Myiasis in filarial lymphoedema due to *Chrysomyia bezziana*. Natl Med J India. 1994;7(3): 117–118.

45. Aguado Lobo M, Hernández-Núnez A, Isabel García-Arata M, Borbujo J. Autochthonous cutaneous myiasis due to *Chrysomya bezziana*. Actas dermo-sifiliograficas. 2014;105(5): 522–524. doi: 10.1016/j.ad.2013.07.005.

46. Center for Health Protection. A case of human myiasis. Communicable disease watch. 2012;9(15): 58.

47. Francesconi F, Lupi O. Myiasis. Clin Microbiol Rev. 2012;25(1): 79–105. doi: 10.1128/CMR.00010-11.

48. Spradbery JP, Kirk J. Incidence of Old World screw-worm fly in the United Arab Emirates. The Vet Rec. 1992;130(2): 33. doi: 10.1136/vr.130.2.33.

49. Sukontason KL, Piangjai S, Boonsriwong, Bunchu N, Ngern-klun R, Vogtsberger RC, Sukontason K. Observations of the third instar larva and puparium of *Chrysomya bezziana* (Diptera: Calliphoridae). Parasitol Res. 2006;99: 699. doi: 10.1007/s00436-006-0233-9.

50. Gan YX. On the larvae of the Chinese species of the subfamily Chrysomyinae (Dipt. Calliphoridae). Dongwuxue Yanjiu. 1980;1: 179–196.

51. Gan YX. On the Chinese species of the genus *Chrysomya Diptera*, Calliphoridae. Acta Entomologica Sinica. 1958;8(4): 340–350.

52. Jarrett S, Morgan JA, Wlodek BM, Brown GW, Urech R, Green PE, et al. Specific detection of the Old World screw-worm fly, *Chrysomya bezziana*, in bulk fly trap catches using real-time PCR. Med Vet Entomol. 2010;24(3): 227–235. doi: 10.1111/j.1365-2915.2010.00867.x.

53. Morgan JA, Urech R. An improved real-time PCR assay for the detection of Old World screw-worm flies. Acta Tropica. 2014;138: 76–81. doi: 10.1016/j.actatropica.2014.02.015.

54. Sabarigirish K, Nithya V, Saxena S, Dutta A. Nasal myiasis by *Chrysomya bezziana*. Medical Journal, Armed Forces India. 2018;74(1): 82–84.

55. Mircheraghi SF, Mircheraghi SF, Ramezani Awal Riabi H, Parsapour A. Nasal Nosocomial Myiasis Infection Caused by *Chrysomya bezziana* (Diptera: Calliphoridae) Following the Septicemia: A Case Report. Int J Dermatol. 2016;11(2): 284–289.

56. Alhady M, Zabri K, Chua CN. Ophthalmomyiasis from *Chrysomyia bezziana* (screwworm fly). The Medical journal of Malaysia. 2008;63: 269–270.

57. Wang JK. A case report of human nasal myiasis. Zhonghua Er Bi Yan Hou Ke Za Zhi. 1960;8: 124.

58. Wadhwa V, Kharbanda P, Rai S, Uppal B. Urogenital myiasis due to *Chrysomyia bezziana*. Indian journal of medical microbiology. 2006;24(1): 70–71.

59. Sankari LS, Ramakrishnan K. Oral myiasis caused by *Chrysomya bezziana*. J Oral Maxillofac Pathol. 2010;14: 16–18.

60. Talari SA, Yeganeh-Moghadam A, Dehghani R,. *Chrysomya bezziana* infestation. Archives of Iranian Medicine. 2002;5(1): 56–58.

61. Ng KH, Yip KT, Choi CH, Yeung KH, Auyeung TW, Tsang AC, et al. A case of oral myiasis due to *Chrysomya bezziana*. Hong Kong Med J. 2003;9(6): 454–456.

62. Nene AS, Mishra A, Dhand P. Ocular myiasis caused by *Chrysomya bezziana* - a case report. Clinical Ophthalmology. 2015;9: 423–427. doi: 10.2147/OPTH.S79754.

63. Lui P, Lee M, Wong J, Leung CY, Lee C, lai R, et al. Myiasis by *Chrysomya bezziana* in surgical pathology. Pathology. 2005;37(1): 80–82. doi: 10.1080/00313020400024758.

64. Ahamd SA, Khan S. Periodontal myiasis treated by open flap debridement: A case report. International journal of medicine and medical sciences. 2012;4(3): 45–48.

65. Souza ND, Kamat S, Chalakkal P, Costa G. A rare occurrence of oral myiasis in the posterior region of the jaw. International Journal of Contemporary Medical Research. 2018;5(8): H6–H8. doi: 10.21276/ijcmr.2018.5.8.11.

66. Kulkarni S, Joshi S, Bhalerao, Chopde Y, Somalwar S. Myiasis: a boon or a bane? J South Asian Feder Obst Gynae. 2012;4(2): 116–117.

67. Prasanna Kumar S, Ravikumar A, Somu L, Vijaya Prabhu P, Mundakannan Subbaiya Periyasamy Subbaraj R. Tracheostomal myiasis: a case report and review of the literature. Case Reports in Otolaryngology. 2011;2011: 303510. doi: 10.1155/2011/303510.

68. Amandeep S, Devinder S. Traumatic myiasis of the scalp in a 3-year-old patient- a case of neglected health care. Malaysian Journal of Medicine & Health Sciences. 2019;15(1): 89–91.

69. Bayer HG. Myiasis maligna of nose and ears in Ceylon recommendation of a new treatment. A M A Archives of otol. 1951;59: 104–107.

70. Zachariah JE, Sehgal K, Dixit UB, Bhatia R. Oral myiasis: a case report. Special Care in Dentistry. 2014;34(1): 51–53.

71. Kalamkar C, Radke N, Mukherjee A. Orbital myiasis in eviscerated socket and review of literature. BMJ Case Reports. 2016;2016: bcr2016215361. doi: 10.1136/bcr-2016-215361.

72. Sharma A, Hedge A. Primary oral myiasis due to *Chrysomya bezziana* treated with Ivermectin, a case report. J Clin Pediatr Dent. 2010;34(3): 259–261.

73. Jimson S, Prakash CA, Balachandran C, Raman M. Oral myiasis: case report. Indian J Dent Res. 2013;24: 750–752.

74. Kumar GV, Sowmya G, Shivananda S. *Chrysomya bezziana* oral myiasis. Journal of global infectious diseases. 2011;3: 393–395.

75. Demograogic Statistics Section(1), Census and Statistics Department. Hong Kong Population Projections (2017-2066). Hong Kong: Census and Statistics Department, Hong Kong Special Administrative Region; 2017.

76. Spradbery JP, Sands DPA, Tozer RS. Distribution and pest status of the Old World screw-worm fly, *Chrysomya bezziana*, in Papua New Guinea and the threat to Australian livestock and wildlife. Austral Entomology. 2019; submitted.

77. Rajapaksa N, Spradbery JP. Occurrence of the Old World screw-worm fly *Chrysomya bezziana* on livestock vessels and commercial aircraft. Aust Vet J. 1989;66(3): 94–96.

78. Sutherst RW, Spradbery JP, Maywald GF. The potential geographical distribution of the Old World screw-worm fly, *Chrysomya bezziana*. Med Vet Entomol. 1989;3: 273–280. doi: 10.1111/j.1365-2915.1989.tb00228.x.

79. Centre for Agriculture and Biosciences International (CABI). Chrysomya bezziana (Old World screw-worm). Avaliable from https://www.cabi.org/isc/datasheet/88417.

80. Hu LL, Ma YH, Chen GG, Guo YF. Myiasis of the female external genitalia caused by *Chrysomya bezziana* in Fujian. Ji Sheng Chong Xue Yu Ji Sheng Chong Bing Za Zhi. 1983;1: 120.

81. Ye YY. Eight cases of myiasis caused by *Chrysomya bezziana* in Guangxi Province. Sheng Chong Xue Yu Ji Sheng Chong Bing Za Zhi. 1985;3: 233.

82. Wu JW, Hu MH, Guo XJ, Tu ZB. A case of mouth myiasis due to *Chrysomya bezziana*. Zhongguo Mei Jie Sheng Wu Xue Ji Kong Zhi Za Zhi. 2008;19: 493.

83. Shi J, Zhao Y, Wang S, Zhang G, Chang M, Liu J, et al. Studies on the family Calliphoridae in Hebei Province (Diptera: Cyclorrhapha). J Med Pest Contrl. 2006;22: 832–835.

84. Fotedar R, Banerjee U, Verma AK. Human cutaneous myiasis due to mixed infestation in a drug addict. Annals of tropical medicine and parasitology. 1991;85: 339–340.

85. Kleine C, Schoefer H, Amendt J, Bexten T, Just-Nuebling G. Cutaneous myiasis in a patient with seborrhoeic eczema. Lancet. 2014;383: 1012. doi:10.1016/S0140-6736(14)60199-9.

86. Rohela M, Jamaiah I, Amir L, Nissapatorn V. A case of auricular myiasis in malaysia. The Southeast Asian journal of tropical medicine and public health. 2006;37(Suppl 3): 91–94.

87. Food and Environmental Hygiene Department. Prevention of myiasis (advice on fly prevention for hospital/homes for the elderly). Hong Kong: Food and Environmental Hygiene Department; 2010.

88. Spradbery JP, Tozer RS, Pound AA. The efficacy of insecticides against the screw-worm fly (*Chrysomya bezziana*). Aust Vet J. 1991;68(10): 338–342.

89. Welch MC, Kwan PW, Sajeev AS. Applying GIS and high performance agent-based simulation for managing an Old World Screwworm fly invasion of Australia. Acta Trop. 2014;138 Suppl: S82–93. doi: 10.1016/j.actatropica.2014.03.021.

90. Lindquist DA, Abusowa M, Hall MJ. The New World screwworm fly in Libya: a review of its introduction and eradication. Med Vet Entomol. 1992;6(1): 2–8.

